# Hypusinated and unhypusinated isoforms of eIF5A exert distinct effects in models of pancreas development and function

**DOI:** 10.1101/2024.10.27.620489

**Authors:** Cara M. Anderson, Abhishek Kulkarni, Bernhard Maier, Fei Huang, Kayla Figatner, Advaita Chakraborty, Sarida Pratuangtham, Sarah C. May, Sarah A. Tersey, Ryan M. Anderson, Raghavendra G. Mirmira

## Abstract

Hypusination of eukaryotic translation initiation factor 5A (eIF5A) is essential for its role in translation elongation and termination. Although the function of hypusinated eIF5A (eIF5A^Hyp^) in cellular proliferation is well-characterized, the role of its unhypusinated form (eIF5A^Lys^) remains unclear. We hypothesized that eIF5A^Lys^ exerts independent effects on cellular replication and metabolism distinct from the loss of eIF5A^Hyp^. To test this hypothesis, we utilized zebrafish and mouse models with inducible knockdowns of deoxyhypusine synthase (DHPS) and eIF5A to investigate their roles in cellular growth. Gene expression analysis via RNA sequencing and morphometric measurements of pancreas and β-cell mass were performed to assess phenotypic changes and identify affected biological pathways. Loss of DHPS in zebrafish resulted in significant defects in pancreatic growth, accompanied by the dysregulation of mRNA translation, neurogenesis, and stress pathways. By contrast, knockdown of eIF5A had minimal impact on pancreas development, suggesting that the effects of DHPS loss are not solely due to the lack of eIF5A^Hyp^. In mice, β cell-specific deletion of DHPS impaired β cell mass expansion and glucose tolerance, while eIF5A deletion had no statistically significant effects. These findings reveal an independent role for eIF5A^Lys^ in regulating developmental and functional responses and that a balance in levels of the hypusinated and unhypusinated isoforms of eIF5A may be pivotal in cellular phenotypes in health and disease.

## Introduction

Protein production is the most energy-consuming cellular process (1). It begins with gene transcription in the nucleus, is followed by translation of the resultant transcript in the cytoplasm, and ends with appropriate folding and posttranslational modifications to generate the final active protein. The process of mRNA translation involves distinct phases of initiation, elongation, and termination, and specific translation factors are known to orchestrate one or more of these phases (see (2) for a review). Regulation of translational rates is crucial to cellular homeostasis, as protein production must respond nimbly to environmental cues and dictate eventual cellular fate, such as replication or apoptosis (3, 4). Translational rates, in turn, are acutely regulated by the posttranslational modification of translation factors or their binding proteins. In states of nutritional surfeit, phosphorylation of 4E binding protein (4EBP) by the mechanistic target of rapamycin (mTOR) releases its inhibition of translation initiation factor 4E (eIF4E), promoting translation initiation and cellular protein production (5). Conversely, during states of amino acid deficiency, the kinase GCN2 phosphorylates translation initiation factor eIF2-α, leading to global suppression of translation initiation and a subsequent reduction in protein synthesis (6, 7). The study of posttranslational modifications and their impact on translation factor function can, therefore, reveal how the environment influences cellular homeostasis.

Eukaryotic translation initiation factor 5A (eIF5A), previously known as eIF4D (8), facilitates the translational elongation and termination of nascent proteins (9), particularly those harboring polyproline residues (10). These functions of eIF5A require an unusual post-translational modification known as hypusine. Hypusine is formed upon the transfer of an aminobutyl moiety from the polyamine spermidine to the epsilon-amino group of Lys50 (11, 12); to date, no other proteins (except the closely related factor eIF5A2) are known to harbor this modification (13). Hypusination is catalyzed by the sequential actions of deoxyhypusine synthase (DHPS) and deoxyhypusine hydroxylase (DOHH), with the former enzyme catalyzing the rate-limiting step (see (14) for review). The balance between unhypusinated eIF5A (eIF5A^Lys^) and hypusinated eIF5A (eIF5A^Hyp^) in a given cell has not been clearly defined, although some prior studies suggest that both forms co-exist (15–17). This means that a readily available pool of eIF5A^Lys^ could exist and be acutely hypusinated in response to environmental cues that demand new protein synthesis. Our prior studies indirectly suggest this possibility, as inducible deletion of *Dhps* in mouse β cells or macrophages impairs protein synthesis and replication in response to environmental stimuli (18, 19). Similarly, in zebrafish, the silencing of *dhps* at an early larval stage leads to defects in pancreatic organogenesis owing to loss of cellular replication (20).

Whereas the cellular phenotypes in these and other studies (21–23) have been attributed to the loss of eIF5A^Hyp^ levels, a role for the concordant increase in eIF5A^Lys^ levels has not been considered. The primary aim of the present study was to elucidate whether eIF5A^Lys^ plays an independent role in previously observed cellular replication phenotypes. In mice, whole-body deletion of *Dhps* or *Eif5a1* (24) or replacement of the endogenous *Eif5a1* alleles with a gene encoding a mutant eIF5A(K50R) (a hypusine modification-incapable mutant) (25) results in embryonic lethality, thereby precluding the use of these models to investigate a role in vivo for eIF5A^Lys^. Instead, we leveraged inducible, timed deletions of the genes encoding DHPS and eIF5A in zebrafish and mouse models to infer roles of eIF5A^Lys^ and eIF5A^Hyp^ by following organismal and cellular phenotypes. Our findings implicate an independent role for eIF5A^Lys^.

## Results

### The loss of total eIF5A does not phenocopy the loss of DHPS

The known roles of eIF5A in mRNA translation are carried out by its active, hypusinated isoform, eIF5A^hyp^ (see Caraglia et al., 2001 for review). Since eIF5A^Lys^ is the only known substrate for the enzyme DHPS, it is generally assumed that loss of the gene encoding eIF5A should largely phenocopy the loss of the gene encoding DHPS. Previous studies showed a growth defect in the exocrine pancreas of 3 days post-fertilization (dpf) zebrafish embryos injected with a splice-blocking morpholino (MO) targeted against *dhps* at the zygote stage (17). To replicate these findings, we utilized the same *dhps* MO, which was designed to be situated at the junction between exon 2 and intron 2 (Fig. 1A). Successful splice blocking should result in two alternatively spliced isoforms of the *dhps* mRNA (**Fig. 1A**). A zygotic injection of 4 ng of *dhps MO* was confirmed by RT-PCR to result in the expected alternatively spliced mRNAs, whereas a control MO resulted in a customarily spliced product (**Fig. 1B**). Also, as expected, no alterations were observed in the levels of total eIF5A in these embryos by immunoblot (**Fig. 1C**). In accord with the prior studies (26), injection of *dhps MO* resulted in a 44% shorter exocrine pancreas at 3 dpf on average, as compared to control MO-injected embryos (**Fig. 1D**). To test if diminished eIF5A protein resulted in a similar short pancreas phenotype, zebrafish zygotes were injected with splice-blocking MOs that targeted each of the zebrafish paralogues, *eif5a1* and *eif5a2* (**Fig. 1E**). Whereas injection of either MO alone did not reduce levels of total eIF5A protein, the combination of both morpholinos (*eif5A1/2* MO) successfully reduced total eIF5A protein (**Fig. 1F**). Despite this reduction in eIF5A protein, we did not observe a significant growth defect in the exocrine pancreas of these embryos (**Fig. 1D**). Together, these results indicate that consequences of DHPS and eIF5A loss are not identical.

**Figure 1:**
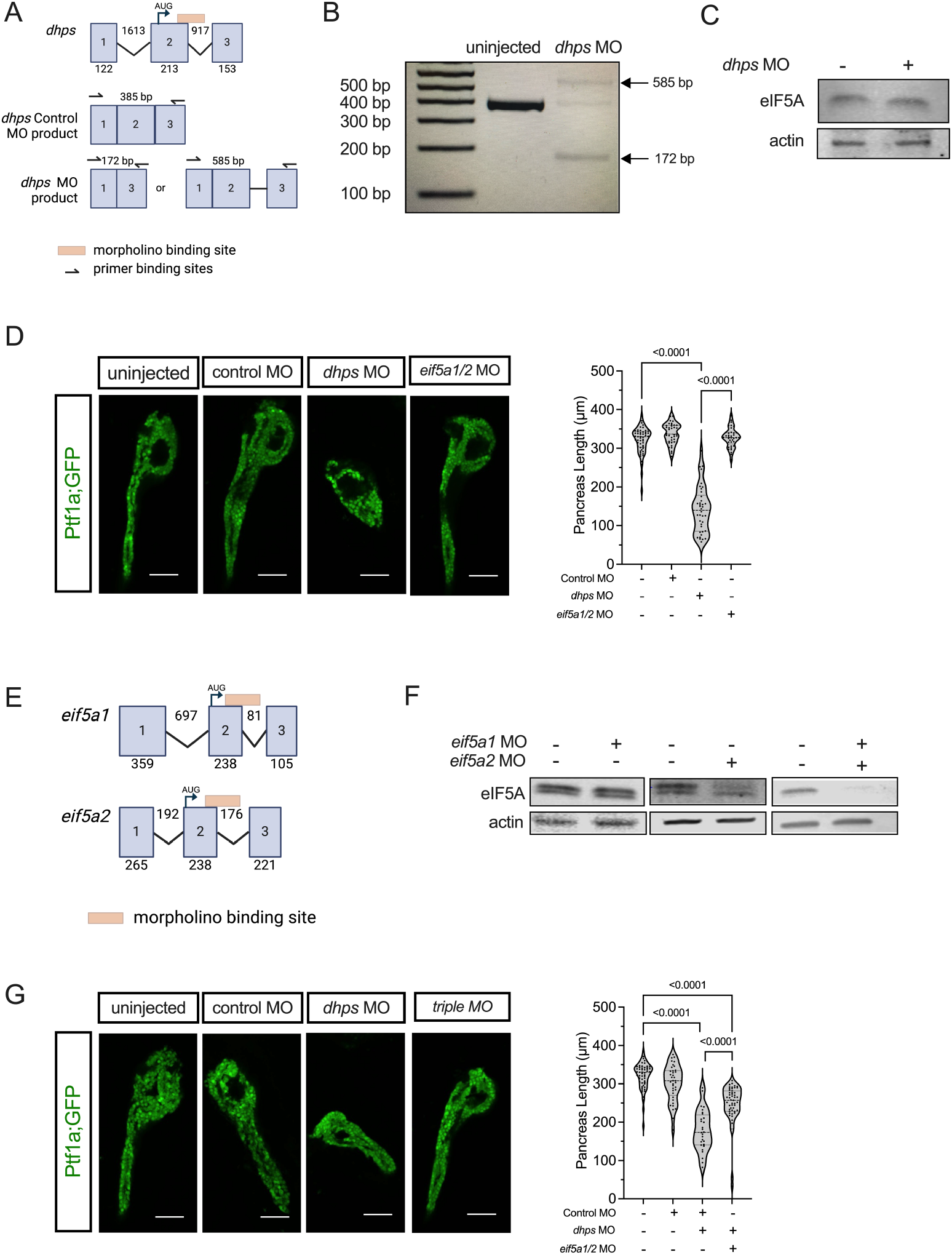
eIF5A^Lys^ is required for phenotypic consequences of DHPS knockdown in zebrafish. (***A***) Model of the first three exons of *dhps* and the binding site of the *dhps* morpholino (MO). PCR products amplified by primers partway through exon 1 (forward) and exon 3 (reverse) are shown: the normally-spliced product and the disrupted-splicing products resulting from 4 ng/embryo *dhps* MO. (***B***) RT-PCR gel showing disrupted *dhps* transcript splicing with *dhps* MO in 24 hours post-fertilization embryos. (***C***) Western blot of total eIF5A protein levels in 0 and 4 ng/embryo *dhps* MO at 3 days post-fertilization (dpf). (***D***) Representative images and quantification of embryonic *Ptf1a;GFP* zebrafish pancreas length at 3 dpf following control, *dhps*, or *eif5a1/2* MO injection. Control MO injected at 8 ng/embryo. *dhps* MO injected at 4 ng/embryo alongside 4 ng/embryo control MO. *eif5a1* MO injected at 4 ng/embryo alongside 4 ng/embryo *eif5a2* MO. Scale bar = 50 μm. Each point on the plot represents one embryo. (***E***) Model of the first three exons and the binding site of the of *eif5a1* and *eif5a2* MOs. (***F***) Western blot of total eIF5A protein knockdown in zebrafish embryos at 3 dpf with a combination of both *eif5a1* and *eif5a2* MOs. Each MO was injected at 4 ng/embryo. (***G***) Representative images and quantification of embryonic *Ptf1a;GFP* zebrafish pancreas length at 3 dpf following control, *dhps*, or combined *dhps* and *eif5a1/2* MO (triple MO) injection. Scale bar = 50 microns. Each point on the plot represents one embryo. Control MO-injected at 12 ng/embryo. *dhps* MO injected at 4 ng/embryo alongside 8 ng/embryo control MO. *dhps, eif5a1*, and *eif5a2* MOs were each injected at 4 ng/embryo together in triple MO embryos. Data are presented as mean ± SEM and statistical significance was determined by a one-way ANOVA with Tukey’s post-hoc test.

### Loss of total eIF5A partially rescues the pancreas phenotype resulting from the loss of DHPS

The silencing of *dhps* should result not only in the loss of eIF5A^hyp^, but also the conjugate accumulation of unhypusinated eIF5A^Lys^; either of these conditions could underlie the exocrine pancreas growth defect. To distinguish between these possibilities, we simultaneously silenced *dhps* and *eif5a1/2* using all three MOs (*triple* MO). If the short pancreas is due simply to the loss of eIF5A^Hyp^ isoform, then the *triple* MO-injected embryos should phenocopy the *dhps* MO-injected embryos. Alternatively, if the short pancreas is due to excess eIF5A^Lys^, then the pancreas of *triple* MO-injected embryos should show a normal or near-normal growth phenotype similar to un-injected embryos. At 3 dpf, we found that the pancreases of *triple* MO-injected embryos were significantly longer than *dhps* MO-injected embryos, though still statistically shorter than those of controls (**Fig. 1G**, *triple* MO vs. *dhps* MO pancreas images). Taken together, these data indicate that eIF5A^Lys^ accumulation likely drives the exocrine pancreas growth defect caused by *dhps* morpholino injection.

### Divergent gene regulatory responses to silencing of dhps versus eif5a1/2 in zebrafish

Although eIF5A is a translation factor, we surmised that its effects on protein production should have secondary consequences on gene expression. These changes would provide insight into the underlying regulatory mechanisms. We, therefore, asked how gene expression is altered following the silencing of *dhps* (leading to accumulation of eIF5A^Lys^) compared to the silencing of *eif5a1/2* (leading to the loss of total eIF5A). Zebrafish zygotes were injected with control MO, *dhps* MO, or *eif5a1/2* MO, then embryos were harvested at 3 dpf and total RNA was subjected to RNA deep sequencing. Principal component analysis of gene expression alterations revealed that the effects of *dhps* MO segregated from controls largely along the component 2 axis (**Fig. 2A**, dotted circles). By contrast, no clear segregation was observed with *eif5a1/2* MO compared to control (**Fig. 2A**). Consistent with the principal component analysis, 1158 genes were altered following *dhps* MO (fold-change, FC, >2 and p-adj <0.05) (**Fig. 2B** and **Supplemental Table 1**). In contrast, fewer than 48 genes were altered following *eif5a1/2* MO (**Fig. 2C** and **Supplemental Table 1**). Of the 48 genes altered with *eif5a1/2* MO, 20 overlapped with the genes altered by *dhps* MO (**Fig. 2D**). Notably, these genes were not changed in the same direction (**Fig. 2E**), suggesting differences in how DHPS and eIF5A individually contribute to distinct cellular pathways (hypusination-dependent or independent). Several of these genes suggest that the two proteins are particularly important in pathways involving protein folding/ER stress (*atf4b, hsp90a*) and cellular apoptosis/proliferation (*casp8, akt1s1, tp53, rspo1, phlda3*).

**Figure 2:**
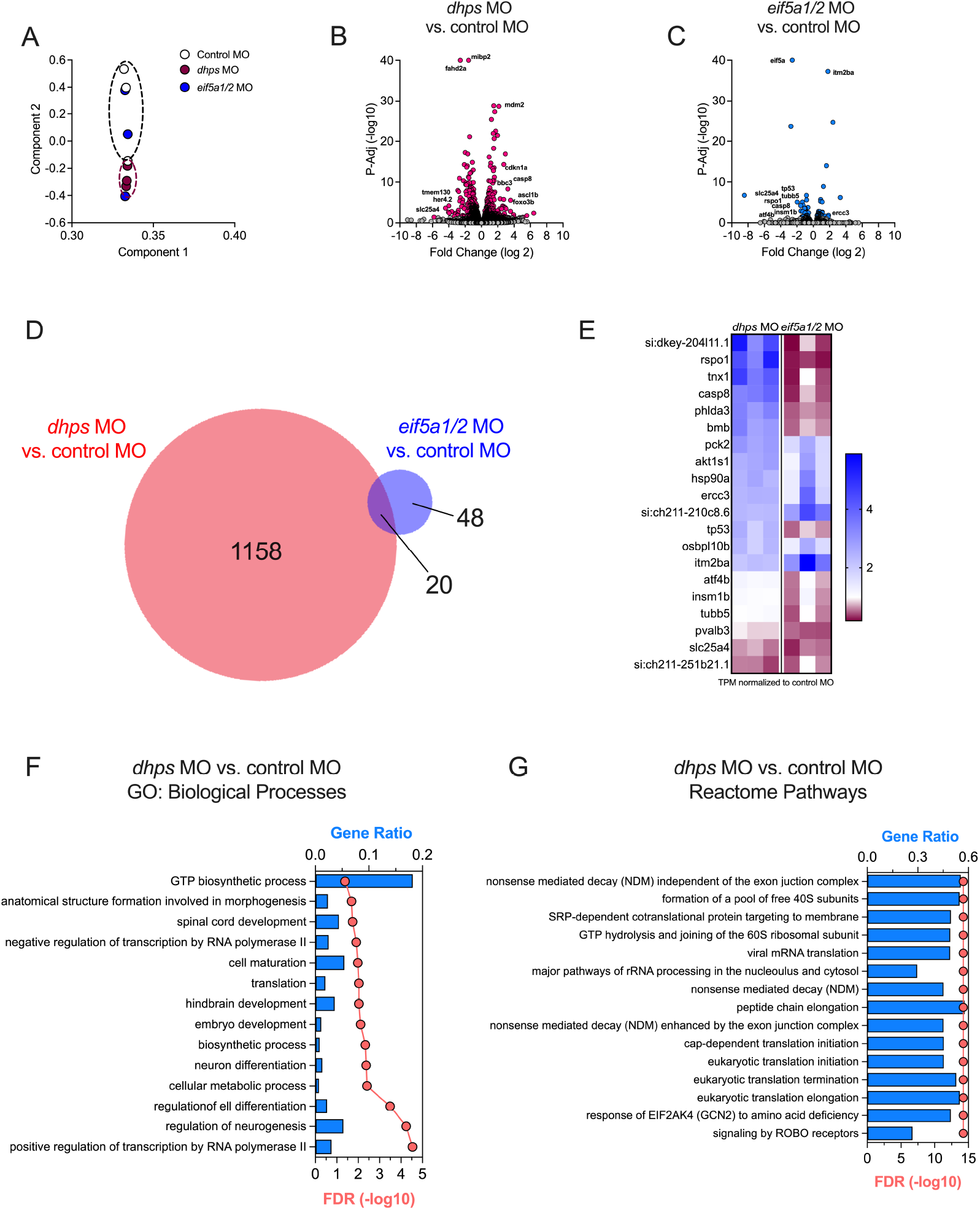
DHPS loss results in different gene expression response compared to eIF5A loss in zebrafish. RNA-sequencing was performed on RAN isolated from zebrafish. Control MO was injected at 8 ng/embryo. *dhps* MO was injected at 4 ng/embryo alongside 4 ng/embryo of control MO. *eif5a1/2* MO denotes embryos injected with 4 ng each of *eif5a1* MO and *eif5a2* MO. (***A***) Principal component analysis of altered gene expression with *dhps* or *eif5a* MO in 3 dpf zebrafish embryos. (***B***) Volcano plot of altered genes in *dhps* MO embryos compared to control MO embryos. (***C***) Volcano plot of altered genes in *eif5a* MO embryos compared to control MO embryos. (***D***) Venn diagram of the overlap between genes altered by *dhps* MO vs. control MO and genes altered by *eif5a* MO vs. control MO. (***E***) Heat map showing the fold change of the 20 genes altered in both *dhps* MO vs. control MO and *eif5a* MO vs. control MO. (***F***) Gene ontology (GO) pathways altered by *dhps* MO vs. control MO. (***G***) Reactome pathways altered by *dhps* MO vs. control MO.

Next, we examined the genes altered explicitly by the *dhps* MO. Genes were clustered based on functional relationships using both Gene Ontology (GO) (**Fig. 2F**) and Reactome (**Fig. 2G**) pathway analyses. The analyses reveal significant disruptions in key biological processes and pathways. Notably, Reactome processes related to mRNA translation, including eukaryotic translation initiation, elongation, and termination, are heavily impacted, indicating a broad effect on protein synthesis. Additionally, pathways like nonsense-mediated decay suggest a role for DHPS in mRNA surveillance mechanisms. GO terms highlight disruptions in developmental processes such as spinal cord and hindbrain development, neuron differentiation, and cellular metabolic processes, reflecting the role of DHPS in cellular growth, differentiation, and metabolic regulation. These findings underscore how the loss of DHPS affects translation and developmental pathways, consistent with the short pancreas phenotype observed with the *dhps* MO.

### Effects of pancreatic β cell deletion of Dhps and Eif5a1 on metabolic phenotypes in mice

We asked if independent effects of DHPS and eIF5A might be observed in mammalian models. Our previous work has shown that pancreatic β cell-specific deletion of *Dhps* in mice leads to impaired β cell replication in response to high fat diet (HFD) feeding (19). However, it was not determined whether this effect was due to the lack of eIF5A^Hyp^ or to the accumulation of eIF5A^Lys^. To distinguish these possibilities, we examined knockouts of *Dhps* and *Eif5a1* in mice. Because whole-body knockout of *Eif5a1* or *Dhps* results in mouse embryonic lethality (24, 27), we used a tamoxifen-inducible Cre recombinase system to temporally and spatially activate the knockouts. We previously described the generation of β cell-specific knockout *Dhps* mice (*Dhps-*Δ*β* mice) (19). We utilized *Eif5a1*^*loxP/loxP*^ mice that we described and validated previously (28) and crossed them to *MIP1-CreERT* mice (29) to produce *Eif5a1*^*loxP/loxP*^; *MIP1-CreERT* mice; upon tamoxifen administration, these mice lack *Eif5a* specifically in the β cell (*Eif5a1-*Δ*β* mice). We induced Cre recombination in β cells between 8-9 weeks of age by tamoxifen administration and followed mice for 5 weeks thereafter (see **Fig. 3A** schematic). *Eif5a1-Δβ* mice had body weight (**Fig. 3B**), body fat mass (**Fig. 3C**), body lean mass (**Fig. 3D**), and glucose tolerance (**Fig. 3E and F**) at 14 weeks of age (5 weeks following the last tamoxifen injection) that were indistinguishable from littermate controls (a combination of *MIP1-CreERT* and *Eif5a1*^*loxP/loxP*^ mice). These data indicate that the post-developmental loss of total eIF5A in β cells has no short-term consequences on metabolic parameters.

**Figure 3:**
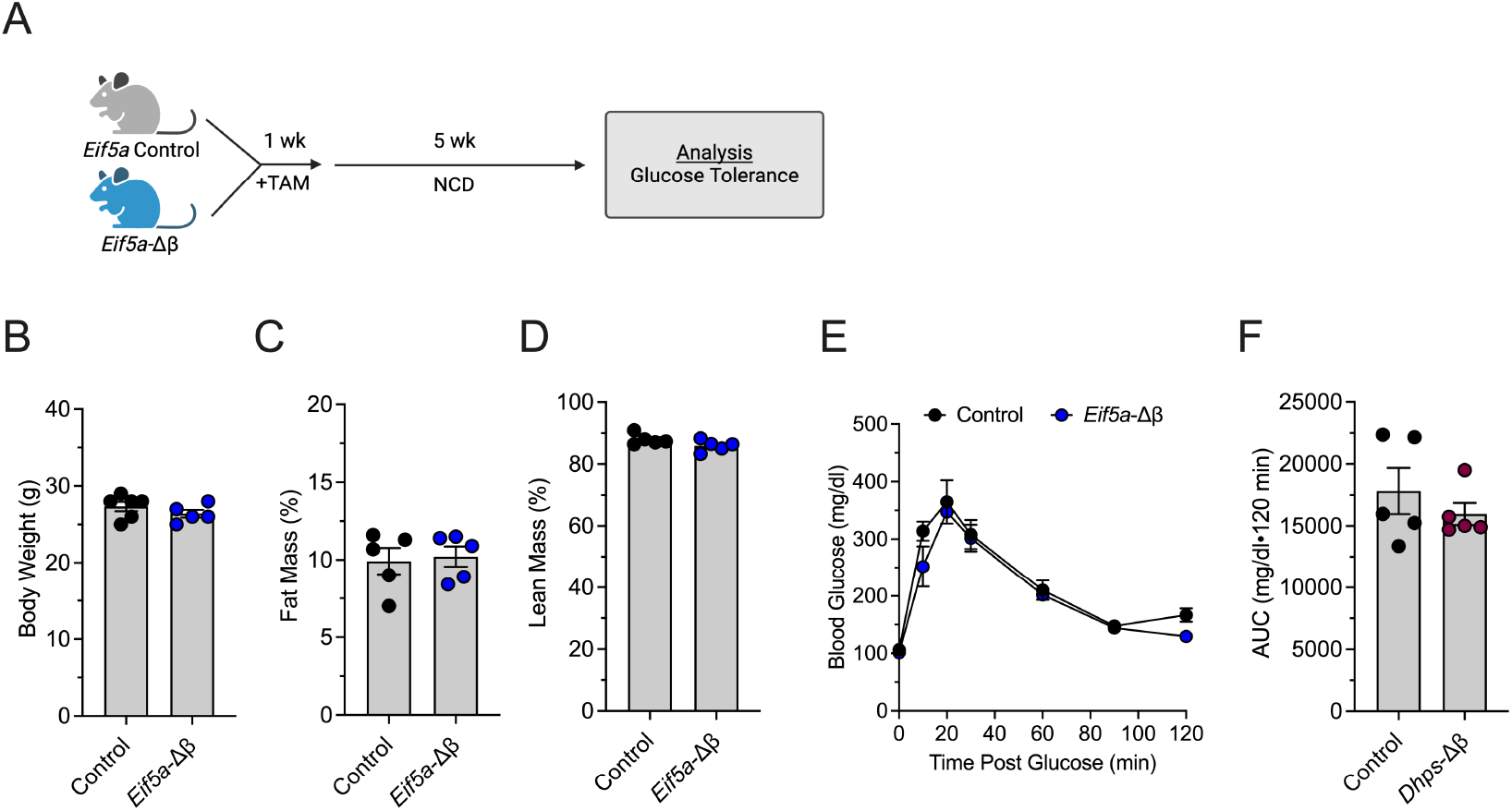
β-cell specific deletion of *Eif5a* in adult mice does not affect glucose homeostasis. 6 week old male *Eif5a*-Δβ and control mice received tamoxifen intraperitoneal injections over one week and allowed a week for recombination. Mice were fed a normal chow diet (NCD) for 5 weeks. (***A***) Diagram showing the experimental design. (***B***) Body weight. (***D***) Fat mass. (***E***) Lean mass. (***F***) Intraperitoneal glucose tolerance test of control and *Eif5a*-Δβ mice. (***G***) Area under the curve (AUC) representation of IPGTT in (***F***). Data are presented as mean ± SEM.

Next, *Eif5a1-Δβ* and *Dhps-Δβ* mice were fed a HFD (60% kcal from fat) for 4 weeks (see **Fig. 4A** for feeding scheme). After 1 week of HFD, *Dhps*-Δβ mice had similar body weights and fat/lean mass compared to their littermate controls (*MIP1-CreERT* and *Dhps*^*loxP/loxP*^) (**Fig. 4B**) and improved glucose tolerance compared to control mice (**Fig. 4C**), consistent with data published previously (19). Likewise, after 1 week of HFD, *Eif5a1*-*Δβ* mice had similar body weights and fat/lean mass compared to their littermate controls (**Fig. 4D**). However, unlike *Dhps-*Δ*β* mice, *Eif5a1-*Δ*β* mice exhibited glucose tolerance comparable to control mice (**Fig. 4E**). After 4 weeks of HFD, *Dhps-Δβ* mice had increased body weight and fat mass that was similar to their littermate controls (**Fig. 4F**); however, at this time point, *Dhps-Δβ* mice exhibited worse glucose intolerance compared to littermate controls (**Fig. 4G**), and similar to previously published data (19). *Eif5a*-Δβ mice had a slight decrease in body weight at 4 weeks, with similar fat and lean mass compared to their littermate controls after 4 weeks of HFD (**Fig. 4H**). Unlike with *Dhps*-Δβ mice, *Eif5a*-Δβ mice exhibited very slightly worse, but statistically insignificant, glucose tolerance compared to controls (**Fig. 4I**). Taken together, these metabolic data are consistent with the pancreatic phenotypic data in zebrafish, where the loss of DHPS (and the presumed increase in eIF5A^Lys^) results in a significantly more dramatic phenotype than the loss of total eIF5A.

**Figure 4:**
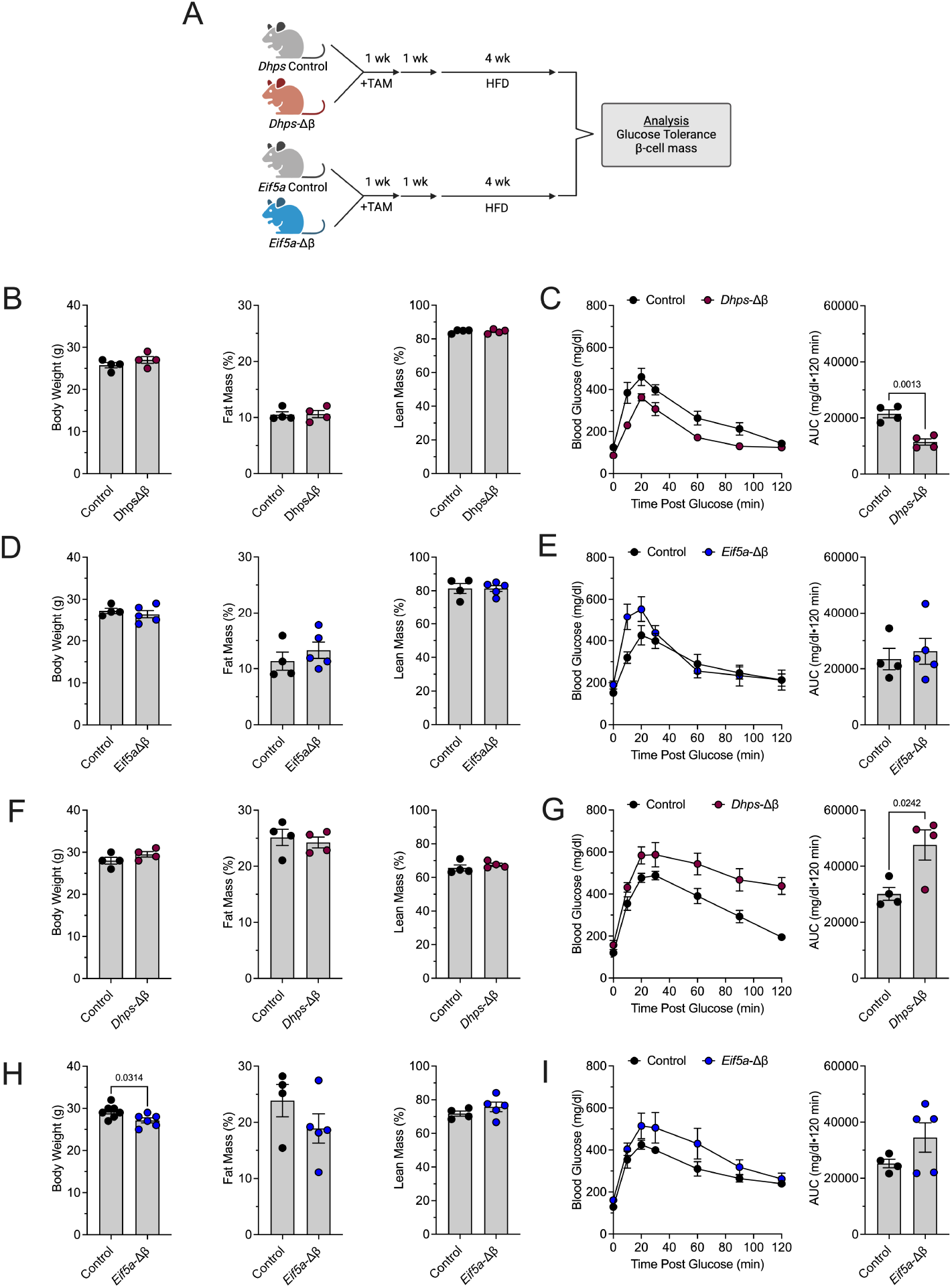
High fat diet induces glucose intolerance in adult *Dhps*-Δβ mice, but not in adult *Eif5a*-Δβ mice. 6-week old male *Eif5a-Δβ, Dhps-Δβ* and their respective control mice received tamoxifen intraperitoneal injections over one week, allowed a week for recombination, and placed on a high-fat diet (HFD, 60% kcal from fat). (***A***) Diagram showing the experimental method of mouse studies. (***B***) Body weight, fat mass, and lean mass of *Dhps-Δβ* mice and *Dhps* control mice after 1 week HFD. (***C***) IPGTT and AUC of *Dhps-Δβ* mice and *Dhps* control mice after 1 week HFD. (***D***) Body weight, fat mass, and lean mass of *Eif5a-Δβ* mice and *Eif5a* control mice after 1 week HFD. (***E***) IPGTT and AUC representation of *Eif5a-Δβ* mice and *Eif5a* control mice after 1 week HFD. Each point represents one mouse. (***F***) Body weight, fat mass, and lean mass of *Dhps-Δβ* mice and *Dhps* control mice after 4 weeks HFD. (***G***) IPGTT and AUC of *Dhps-*Δβ mice and *Dhps* control mice after 4 weeks HFD. (***H***) Body weight, fat mass, and lean mass of *Eif5a-Δβ* mice and *Eif5a* control mice after 4 weeks HFD. (***I***) IPGTT and AUC of *Eif5a-Δβ* mice and *Eif5a* control mice after 4 weeks HFD. Data are presented as mean ± SEM and statistical significance was determined by an unpaired t-test.

### Loss of Dhps, but not Eif5a1, reduces β cell mass and affects proliferation and stress responses

The effects on glucose tolerance following HFD feeding of *Dhps-*Δ*β*, and *Eif5a1-*Δ*β* mice led us to investigate how β cells might be affected in these animals. We isolated pancreata from mice following 4 weeks of HFD feeding and performed a morphometric analysis of β cell mass. Consistent with our prior study (19), we found a statistically reduced β cell mass in the *Dhps-Δβ* mice compared to their littermate controls (**Fig. 5A**), suggesting that the worsened glucose tolerance in these animals might be attributed to the loss of β cell insulin compensation. By contrast, *Eif5a1-Δβ* mice exhibited β cell mass similar to their littermate controls (**Fig. 5A**). This loss in cellular mass in HFD-fed *Dhps-*Δ*β* mice (and contrasting preservation in HFD-fed *Eif5a1-*Δ*β* mice) is reminiscent of the loss in pancreas length that we observed in *dhps* MO zebrafish (and preservation seen in *eif5a1/2* MO zebrafish). To determine if similar gene response pathways are observed in HFD-fed *Dhps-*Δ*β* islets, we re-analyzed islet RNA sequencing data from our prior study of 4-week HFD-fed *Dhps-*Δ*β* mice and littermate controls (19). Using a cut-off of FC>2 and p-adj <0.05, there were 1619 differentially expressed genes in *Dhps-*Δ*β* mice compared to controls (**Fig. 5B** and **Supplemental Table 2**).

**Figure 5:**
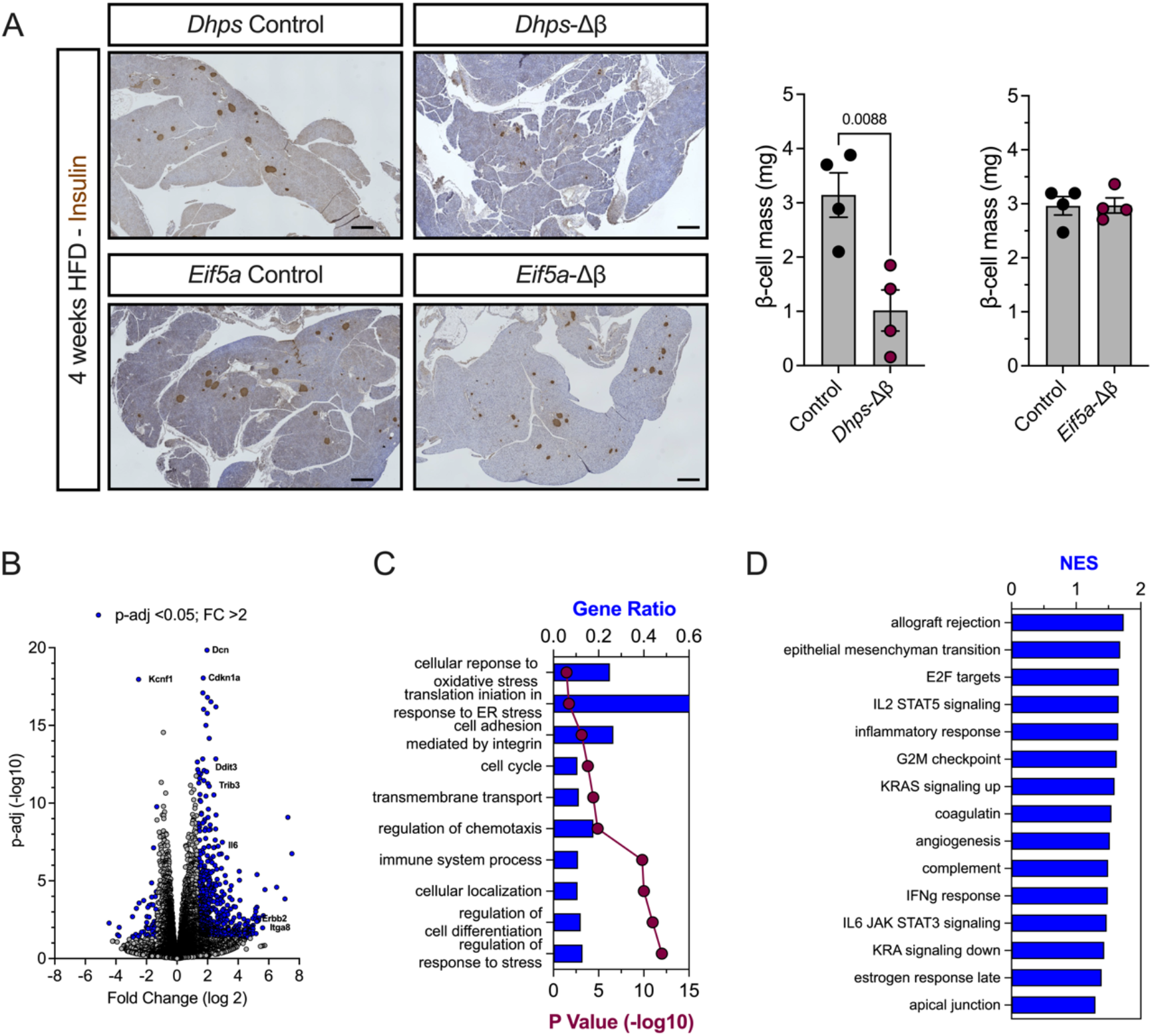
*Dhps* deletion, but not *Eif5a* deletion, in adult mouse β cells impairs β cell mass expansion in response to 4 weeks HFD. (***A***) Representative images and quantification of insulin-positive β cell mass in *Dhps* control, *Dhps-Δβ, Eif5a* control, and *Eif5a-Δβ* mouse pancreases after 4 weeks HFD. Scale bar = 100 μm. Each point represents one mouse. (***B***) Volcano plot showing genes with altered expression in *Dhps-Δβ* mouse islets compared to *Dhps* control mouse islets. (***C***) Gene ontology (GO) pathways of *Dhps-Δβ* mouse islets compared to *Dhps* control mouse islets. (***D***) Gene set enrichment analysis of *Dhps-Δβ* mouse islets compared to *Dhps* control mouse islets. Data are presented as mean ± SEM and statistical significance was determined by an unpaired t-test.

The GO pathway analysis (**Fig. 5C**) and gene set enrichment analyses (GSEA) (**Fig. 5D**, depicting normalized enrichment scores) from *Dhps-*Δ*β* islets reveal involvement in pathways related to oxidative stress response, ER stress response, cell cycle regulation, and immune system processes. This finding suggests that *Dhps* knockout in β cells affects cellular stress responses, immune signaling, and proliferation mechanisms. Comparing these findings to the GO analysis from the zebrafish *dhps* MO, both highlight translation-related processes, particularly the response to cellular stress and developmental signaling pathways. In zebrafish, the focus was more on translation and developmental processes like neurogenesis and morphogenesis, while in β cells, immune signaling and cell adhesion pathways are more prominent. These findings suggest a conserved role for DHPS in regulating cellular stress and protein synthesis, with context-specific effects on immune and developmental processes depending on the tissue type.

## Discussion

It has long been appreciated that DHPS and eIF5A hypusination are essential to cellular replication (19, 30–34). This and other physiological consequences of DHPS inhibition have been ascribed mainly to the reduced abundance of eIF5A^Hyp^, due to the well-documented role of eIF5A^Hyp^ in facilitating translation elongation and termination (9, 35–38). However, inhibiting DHPS also results in an increase in eIF5A^Lys^, whose function has largely been understudied. To date, there is a lack of evidence that eIF5A^Lys^ has a cellular function independent of the translation elongation role of eIF5A^Hyp^. In this study, we used zebrafish and mouse models to interrogate how the loss of DHPS (and the consequent accumulation of eIF5A^Lys^) compares to the loss of total eIF5A (both eIF5A^Lys^ and eIF5A^Hyp^). Our findings suggest that acutely, the loss of DHPS is more severe, suggesting that an accumulation of eIF5A^Lys^ may have an independent effect from the loss of eIF5A altogether.

Hypusine is an unusual spermidine-derived amino acid that is post-translationally formed within eIF5A in a two-step reaction, wherein the lysine at position 50 of eIF5A is converted first to deoxyhypusine by DHPS, then to hypusine by DOHH (12, 14). Inhibiting or deleting the gene encoding DHPS results in stark cellular phenotypes, including impaired cell replication, growth, and function, attributed to the decrease in eIF5A^Hyp^ (19–21, 39). In addition, neurodevelopmental disorders in humans, mice, and zebrafish have been attributed to the loss or dysfunction of DHPS (40–43). By comparing the consequences of DHPS loss with total eIF5A loss side-by-side in this study, we are able to comment on the effects of accumulated eIF5A^Lys^ compared to the loss of total eIF5A. In zebrafish, *dhps* MO resulted in a stunted pancreas growth phenotype in the short term. Conversely, *eif5a1/2* MO resulted in normal pancreas morphogenesis during the same time frame. At first glance, these data seemingly contrast with previous studies showing that loss of total eIF5A suppresses cell replication to a greater degree than the loss of DOHH analog nero in *D. melanogaster* (44). However, it is possible that the accumulation of deoxyhypusinated eIF5A accompanying the loss of DOHH in that study mitigated an otherwise more severe phenotype that could have occurred with a loss in DHPS. It should be noted that another study that examined the loss of total eIF5A alongside loss of DHPS reported more severe phenotypes with the latter (41). In our study, the partial rescue of pancreas length by combined *dhps* MO and *eif5a1/2* MO (*triple* MO) refutes the notion that the DHPS-loss phenotype is driven entirely by lack of eIF5A^Hyp^. The DHPS-loss phenotype is mitigated by reduction of total eIF5A in zebrafish embryos, indicating that the presence eIF5A^Lys^ appears to have, at least acutely, an independent role.

Our RNA sequencing data in zebrafish provide new insights into the underlying mechanisms driving phenotypes in *dhps* MO and *eif5a1/2* MO zebrafish. Several key features arise from our analysis: (a) gene expression changes can occur with factors traditionally associated with the translational machinery, reflecting the molecular crosstalk between translation and transcription in the cell, (b) the number of genes altered acutely by *dhps* MO greatly exceed those altered by *eif5a1/2* MO, correlating with the more pronounced morphological phenotypes observed with *dhps* silencing, (c) the genes altered by *dhps* MO were enriched in pathways that affect cellular differentiation, replication, and mRNA translation, suggesting that the loss of DHPS, unlike eIF5A, acutely drives dysmorphogenesis, and (d) genes that are common following silencing of *dhps* and *eif5a1/2* reflect processes related to mRNA translation/ER stress and cellular proliferation/apoptosis. It is important to note that these studies examined only the short-term effects of *dhps* and *eif5a1/2* silencing (within hours), and longer-term studies may reveal molecular and morphological phenotypes that converge more closely over time (41, 42).

Isolating the physiological consequences of total eIF5A or DHPS loss in mice has been challenging due to the early embryonic lethality associated with β-cell-specific knockouts of eIF5A or DHPS (24, 27). To circumvent this limitation, we used the *MIP1-CreERT* mouse strain (29) to induce gene knockouts in β cells at 8 weeks of age. In a previous study, we demonstrated that *Dhps* knockout in mature β cells impairs the expansion of β-cell mass in response to a HFD (19). Here, we show that β cells lacking eIF5A do not exhibit this impairment in β-cell mass accrual. Our data suggest that the accumulation of eIF5A^Lys^ following *Dhps* knockdown may actively suppress adaptive β-cell replication. Given that DHPS activity is essential for the proliferation of several tumor types and that eIF5A expression is notably elevated in various malignancies (see (45) for review), further investigation into the specific effects of eIF5A^Lys^ accumulation in other cell types is warranted.

Our study provides an initial exploration into the potential consequences of eIF5A^Lys^ accumulation versus the loss of eIF5A^Hyp^, but several important questions remain. What are the specific mechanisms through which eIF5A^Lys^ might function? Is there a regulated balance between eIF5A^Lys^ and eIF5A^Hyp^ within cells, and how does disruption of this balance affect cellular function, cancer progression, or other disorders? Clues from the literature offer insight. For instance, the structure of eIF5A^Hyp^ has been shown to share three-dimensional homology with that of charged tRNA (46). This raises the possibility that eIF5A^Lys^, like an uncharged tRNA, could activate molecular pathways such as the integrated stress response via GCN2 kinase (see Costa-Mattioli and Walter, 2020 for review). Supporting this idea, our RNA sequencing data from *dhps* silencing in zebrafish revealed enrichment in Reactome pathways, including “eukaryotic translation initiation,” “cap-dependent translation initiation,” and “response of EIF2AK4 (GCN2) to amino acid deficiency”—all of which are associated with the integrated stress response. Collectively, our findings suggest a potential independent role for eIF5A^Lys^, and elucidating its precise function and possible molecular partners represents a key direction for future research.

## Materials and methods

### Zebrafish and mouse strains

The *Tg(ptf1a:eGFP)*^*jh1*^ zebrafish transgenic line (48) was used for all zebrafish experiments. Zebrafish were maintained and inter-bred under standard conditions at 28°C. *C57BL/6J*.*Dhps*^*loxp*^ (19) or *C57BL/6J*.*Eif5a*^*loxp*^ (28) mice were crossed to *C57BL/6J*.*MIP1-CreERT* (29) mice to generate *Dhps-Δβ* mice and *Eif5a-Δβ*, respectively. All mice were bred in-house in specific pathogen-free conditions and maintained on a 12 h light/12 h dark cycle with free access to food and water. All animal studies were performed under protocols approved by the University of Chicago Institutional Animal Care and Use Committee.

### Morpholino design and injection

25-base morpholinos were purchased from GeneTools, LLC, dissolved in water, and injected into zebrafish zygotes prior to the first cell division using a uPUMPmicroinjector (World Precision Instruments). The standard control morpholino (5’– CCTCTTACCTCAGTTACAATTTATA–3’) was an AUG-binding translation-blocking design, while all others were splice-blocking, and each bound at the exon 2-intron 3 junction of the preRNA transcripts: *dhps* morpholino (5’–ACGATCAGTCTGTCACTCACCATCT–3’, 4 ng injected per zygote) (20); *eif5a1* morpholino (5’–AACCCTATCCAAACATTACCTTTGC–3’, 4 ng); *eif5a2* morpholino (5’–TTATTAATACGACACCTTGGCATGT–3’, 4 ng). RT-PCR was used to evaluate the efficacy of *dhps* morpholino as previously described (20). Protein quantification by immunoblot was used to assess the effect on eIF5A expression by the *dhps* morpholino and by combining *eif5a1* and *eif5a2* morpholinos (BD Pharmingen).

### Pancreas length quantification

At 3 days post-fertilization, zebrafish embryos were anesthetized with Tricaine and fixed in a 2.5% formaldehyde/PEM solution for 18 hours at 4°C. Then, the embryos were transferred to PBS and microdissected to remove skin and yolk before being dehydrated in 100% methanol. Embryos were rehydrated into PBS + 0.1% Tween-20 and then mounted on a glass microscope slide. Pancreata were visualized with (Nikon A1 confocal microscope) at 40X magnification. Pancreas length was quantified in microns by manual measurement in ImageJ.

### RNA sequencing

Total RNA was harvested from 24 hours post-fertilization zebrafish embryos through Trizol-chloroform extraction. Reads were aligned to the GRCz11 genome build. Sequencing was performed by Innomics and subsequent analysis was conducted using Dr. Tom web-based platform (BGI). Pathways for both the zebrafish RNA-sequencing results and the high fat diet-fed *Dhps*-Δβ mice RNA-sequencing were analyzed or reanalyzed using the Biological Function Gene Ontology (GO) and Reactome Pathway Browser.

### Mouse studies

At 8 weeks of age, *Dhps-Δβ* mice, *Eif5a-Δβ*, and their littermate control mice received intraperitoneal injections of 2.5 mg tamoxifen dissolved in peanut oil once daily for three days. After 1 week of rest, the mice were fed a normal chow diet (6% kcal from fat) (Envigo) or a high-fat diet (60% kcal from fat) (Research Diets) for 4 weeks. After 1 week and 4 weeks on these diets, body mass was measured by EchoMRI, and intraperitoneal glucose tolerance tests were performed based on lean mass as previously described (16). At the end of the study, mice were euthanized, and pancreas was harvested (49).

### Immunostaining, immunoblotting, and β cell mass measurements

For immunoblotting of whole zebrafish embryos, approximately 50 embryos at 3 dpf per condition were anaesthetized with tricaine, then lysed in lysis buffer (Thermo Fisher) and subjected to electrophoresis on a 4%–20% gradient SDS–polyacrylamide gel. Antibodies included mouse monoclonal antibody directed against eIF5A (1:1000; BD Biosciences #611977) and mouse monoclonal antibody against β-actin (1:1000, Cell Signaling #3700s). Immunoblots were visualized using fluorescently labeled secondary antibodies (LI-COR Biosciences) and were quantified using LI-COR software. For immunohistochemistry, pancreata were fixed in 4% paraformaldehyde, paraffin-embedded, and sectioned. For β cell mass, pancreas sections were stained with anti-insulin (1:200; ProteinTech #15848-1-AP) followed by Impress reagent kit peroxidase-conjugated anti-rabbit Ig (Vector Laboratories), DAB peroxidase substrate kit (Vector Laboratories), and counterstained with hematoxylin (Sigma). β cell mass was measured in 3 sections at least 100 μm apart and images were collected on a BZ-X810 (Keyence). β cell mass was calculated as a ratio of the insulin+ area and whole pancreas area using the BZ-X810 Analyzer (Keyence).

### Statistical analysis

For comparisons between two conditions, a two-tailed unpaired T-test was performed. A one-way ANOVA test with Tukey’s post hoc test was performed for comparisons with more than two conditions. GraphPad Prism was used for all statistical analysis.

## Acknowledgments

This work was supported by NIH grants R01 DK124906 and R01 DK060581 (both to RGM), R01 DK135832 (to SAT), and F31 DK134070 (to CMA). This work utilized core services provided by the Diabetes Research and Training Center award P30 DK020595 (to the University of Chicago). Funders played no part in the design or interpretation of the experiments.

## Author contributions

CMA, BM, SAT, RMA, and RGM designed and conceptualized the research; CMA, AK, FH, KF, AC, and SP performed research; RGM, SAT, and CMA obtained funding for the project; CMA, SCM, SAT, RMA, and RGM wrote the initial draft of the manuscript; All authors edited and approved the final version of the manuscript.

## Data availability

The zebrafish RNA sequencing data have been uploaded to the Gene Expression Omnibus (https://www.ncbi.nlm.nih.gov/geo/) with accession number GSE279773. Any additional information required to reanalyze the data reported in this study is available from the corresponding author upon request.

